# The long-term effects of repeated heroin vapor inhalation during adolescence on measures of nociception and anxiety-like behavior in adult Wistar rats

**DOI:** 10.1101/2021.10.06.463404

**Authors:** Arnold Gutierrez, Eric L. Harvey, Kevin M. Creehan, Michael A. Taffe

## Abstract

**Rationale:** Adolescents represent a vulnerable group due to increased experimentation with illicit substances that is often associated with the adolescent period, and because adolescent drug use can result in long-term effects that differ from those caused by drug use initiated during adulthood.

**Objectives:** The purpose of the present study was to determine the effects of repeated heroin vapor inhalation during adolescence on measures of nociception, and anxiety-like behavior during adulthood in female and male Wistar rats.

**Methods:** Rats were exposed twice daily to 30-minutes of heroin vapor from post-natal day (PND) 36 to PND 45. At 12 weeks of age, baseline thermal nociception was assessed across a range of temperatures with a warm-water tail-withdrawal assay. Anxiety-like behavior was assessed in an elevated plus-maze (EPM) and activity was measured in an open field arena. Starting at 23 weeks of age, baseline thermal nociception was re-assessed, nociception was determined after acute heroin or naloxone injection, and anxiety-like behavior was redetermined in the EPM.

**Results:** Adolescent heroin inhalation altered baseline thermal nociception in female rats at 12 weeks of age and in both female and male rats at ∼23 weeks. Heroin-treated animals exhibited anxiety-like behavior when tested in the elevated plus-maze, showed blunted heroin-induced analgesia, but exhibited no effect on naloxone-induced hyperalgesia.

**Conclusions:** The present study demonstrates that heroin vapor inhalation during adolescence produces behavioral and physiological consequences in rats that persist well into adulthood.

## Introduction

The US opioid crisis, which is associated with high rates of non-medical use of both prescription and illicit opioids, has been marked by escalating rates of opioid-associated overdose and increasing prevalence of addictive use disorders (Han et al. 2020; SAMHSA/CBHSQ 2018; TEDS/SAMHSA 2018; Wilson et al. 2020). Treatment admissions of adolescents and young adults for heroin use is second only to admissions for cannabis in the US (Standeven et al. 2020). At the same time, there has been an alarming rise in the use of Electronic Nicotine Delivery Systems (ENDS; e-cigarettes), especially within adolescent populations. Adolescents who administer nicotine using ENDS devices are more likely to also use them to vape cannabis (Merianos et al. 2019), and, among college students, ENDS devices are increasingly being used for the administration of substances other than nicotine (Kenne et al. 2017). ENDS devices can be readily adapted to deliver a range of psychoactive drugs (Breitbarth et al. 2018), and reports are emerging on their use to administer substances such as cathinones, amphetamines, cocaine, and opioids (Blundell et al. 2018; Thurtle et al. 2017). One such report found that 7.1% of those who reported any vaping experience and 25.8% of those who reported current usage of electronic vapor devices for non-nicotine substances had vaped heroin, with similar numbers (7.3% and 26.7%, respectively) reported for vaped fentanyl (Blundell et al. 2018). The popularity of ENDS for administering non-nicotine substances has reached a level that recommends rethinking the terminology; in truth, Electronic Drug Delivery Systems (EDDS) is a more accurate term.

Adolescence represents a critical developmental period associated with increased risk-taking and impulsive behavior (Spear 2000). Experimentation with drugs and alcohol is heightened during this period, and adolescents are therefore at an increased risk of developing substance abuse disorders (Chambers et al. 2003; Wagner 2002). In animal models, chronic drug exposure during adolescence can result in long-lasting effects on nociception and drug reinforcement. For example, mice that self-administered oxycodone during adolescence exhibit a decrease oxycodone anti-nociception in adulthood (Zhang et al. 2016) and tolerance to morphine anti-nociception is accelerated in adult rats that were chronically treated with morphine during adolescence (Salmanzadeh et al. 2017). Morphine conditioned place preference (CPP) is enhanced in adult mice treated with oxycodone (Sanchez et al. 2016) and in adult rats treated with morphine during adolescence (Schwarz and Bilbo 2013). Similarly, oxycodone self-administration in adolescence potentiates oxycodone CPP in adult mice (Zhang et al. 2016). While the lasting effects of opioid treatment during adolescence have been studied in animals, to date they have mainly been studied using parenteral injections, or implanted delivery systems, as the route of administration. Therefore, the effects of inhaled opioids within a developmental context, if any, are presently unknown.

Inhalation is a common route of administration among opioid users (Alambyan et al. 2018) that dates back to Mediterranean antiquity (Kritikos 1960). The inhalation of opioids can be achieved by heating the drug over a surface such as aluminum foil and inhaling the resulting vapors, a method often referred to as ‘chasing the dragon’ (Jenkins et al. 1994; Strang et al. 1997). Initial studies confirmed anti-nociceptive effects of heroin vapor inhalation, which were reversed with the opioid receptor antagonist naloxone, in mice (Lichtman et al. 1996), and reported self-administration of the opioid sufentanil in aerosol mist form through the use of a nebulizer in rats (Jaffe et al. 1989). Additionally, self-administration of heroin by inhalation was reported in rhesus monkeys (Evans et al. 2003; Foltin and Evans 2001; Mattox and Carroll 1996; Mattox et al. 1997). These initial studies using inhalation as the route of administration provided a translationally relevant alternative approach to the more commonly used parenteral routes of drug administration. This is especially true when implanted delivery systems are considered. Additionally, the vapor model avoids the stress produced by injection, especially when multiple daily injections are involved. Following these initial studies on the topic, a dearth of preclinical literature on the effects of inhaled opioid drugs remained until the recent adaptation of ENDS/EDDS technology for preclinical drug inhalation research.

EDDS technology can be adapted to deliver active doses of many psychoactive drugs to rats by inhalation, including opioids. Non-contingent vapor delivery produces anti-nociception after inhalation of Δ9-tetrahydrocannabinol (THC) vapor (Gutierrez et al. 2022; Javadi-Paydar et al. 2018; Nguyen et al. 2016b), decreases body temperature after nicotine, THC or cannabidiol inhalation (Gutierrez et al. 2022; Javadi-Paydar et al. 2019a; Javadi-Paydar et al. 2019b; Javadi-Paydar et al. 2018; Nguyen et al. 2016b), increases locomotor activity after exposure to nicotine (Javadi-Paydar et al. 2019b), and methamphetamine (Nguyen et al. 2016a; Nguyen et al. 2017), and decreases activity following THC (Gutierrez et al. 2022). Additionally, EDDS delivery of nicotine vapor produces CPP in adult and adolescent rats (Frie et al. 2020). *In utero* exposure to vaporized THC produces effects on body weight in adolescence (Breit et al. 2020), and a recent study shows evidence of the self-administration of THC by vapor inhalation (Freels et al. 2020). Recent studies also present compelling evidence for the inhalation self-administration of nicotine by rats (Lallai et al. 2021; Smith et al. 2020) and mice (Cooper et al. 2021; Henderson and Cooper 2021). Inhalation of oxycodone, methadone, and heroin vapor produces anti-nociception (Gutierrez et al. 2021; Gutierrez et al. 2022; Gutierrez et al. 2020; Nguyen et al. 2019), and heroin vapor inhalation elevates body temperature and produces biphasic effects on activity in rats (Gutierrez et al. 2021). Self-administration of opioids by inhalation using EDDS technology has also recently been reported in rodents. Mice and rats will make responses to obtain doses of fentanyl vapor (McConnell et al. 2020; Moussawi et al. 2020), and rats will make responses to obtain doses of the potent opioid sufentanil in vapor form in an extended daily access paradigm (Vendruscolo et al. 2018) and heroin vapor in a short-access paradigm (Gutierrez et al. 2020). Importantly, signs of dependence were observed in both the mouse and rat opioid vapor self-administration studies described.

Opioid drugs, which are clinically used for the treatment of pain, can produce alterations in nociception, such as opioid tolerance and opioid-induced hyperalgesia (Mao et al. 1995; Prommer 2006), and can result in negative affective symptoms (Kanof et al. 1993; Wermuth et al. 1987). Importantly, alterations in nociception are commonly reported in opioid abusers and are associated with increased drug craving (Ren et al. 2009; Tsui et al. 2016). Similarly, anxiety disorders are associated with an increased risk of developing substance abuse disorders (Bushnell et al. 2019; Lopez et al. 2005). The purpose of this study was to assess the long-term effects of repeated adolescent heroin vapor exposure on thermal nociception, injected opioid anti-nociception, and on anxiety-like behavior in adulthood. Following repeated heroin vapor exposure during adolescence and upon reaching adulthood, rats were tested for differences in nociceptive sensitivity using a warm water tail-withdrawal assay. Anxiety-like behavior was assessed using an elevated plus-maze (EPM) and open field tests. Finally, anti-nociceptive effects of acute heroin injection and hyperalgesic effects of injection with the opioid receptor antagonist naloxone were determined. This is the first study to assess the long-term effects of adolescent opioid exposure using inhalation as the route of administration.

## Methods

### Animals

Adolescent female (N=24) and male (N=24) Wistar rats were obtained from Charles River Laboratories (Charles River) in two sex-balanced cohorts. Two cohorts of female (N=12 per cohort) and male (N=12 per cohort) rats, designated Cohort 1 (C1) and Cohort 2 (C2), arrived on Post-natal Day (PND) 29 and were habituated to the vivarium through PND 35. The vivarium was kept on a 12:12 hour light-dark cycle and behavior studies were conducted during the rats’ dark period. Food and water were provided ad libitum in the home cage. Body weights were recorded beginning immediately prior to vapor drug treatment at around 5 weeks of age (PND 36), one week after commencement of vapor drug treatment at 6 weeks of age (PND 43), and after completion of vapor inhalation sessions on week 7 (PND 51). Importantly, weights during this time were not affected by Adolescent Drug treatment. Body weights were then recorded weekly through the end of the study; these data are reported in the **Supplementary Information (Fig. S1**). Experimental procedures were conducted in accordance with protocols approved by the IACUC of the University of California, San Diego and consistent with recommendations in the NIH Guide (Garber et al. 2011).

Rats were randomly assigned to the experimental (heroin inhalation) or control (propylene glycol inhalation) Adolescent Drug groups. Group sizes were determined based on prior work (Gutierrez et al. 2021; Gutierrez et al. 2022; Gutierrez et al. 2020; Javadi-Paydar et al. 2018; Nguyen et al. 2018) Gutierrez et al., 2020; Gutierrez et al., 2021) and were designed to be equal. However, two male rats from Cohort 1 (C1) assigned to the repeated heroin group were lost during repeated vapor exposure presumably due to group huddling, immobility due to intoxication and perhaps respiratory suppression. Thereafter a procedure was adopted in which the experimenter periodically gently jostled all exposure chambers to prevent static huddles. With the exception of the EPM data, repeated measures designs were employed, thus minimizing the number of animals used while increasing statistical power.

### Drugs

Heroin (diamorphine HCl) was administered by vapor inhalation and subcutaneous injection. Inhalation/vapor doses are varied in this model, and are therefore described, by altering the concentration in the propylene glycol (PG) vehicle, the puff schedule and/or the duration of inhalation sessions in this approach (Javadi-Paydar et al. 2019b; Nguyen et al. 2016a; Nguyen et al. 2016b). The heroin was dissolved in the PG to achieve target concentrations and then loaded in the EDDS tanks in a volume of ∼ 0.5 ml per session. PG was used as the vehicle to enhance consistency and comparison with our prior publications on the topic of heroin vapor inhalation. Fresh solutions were used for each session. Heroin and naloxone (naloxone HCl) were dissolved in physiological saline for subcutaneous (s.c.) or intraperitoneal (i.p.) injection, respectively. Heroin was provided by the U.S. National Institute on Drug Abuse. Naloxone (Tocris Bioscience) and propylene glycol were purchased from Fisher Scientific.

### Vapor drug treatment

Vapor was delivered into sealed vapor exposure chambers (152 mm W X 178 mm H X 330 mm L; La Jolla Alcohol Research, Inc, La Jolla, CA, USA) through the use of e-vape controllers (Model SSV-3 or SVS-200; 58 watts; La Jolla Alcohol Research, Inc, La Jolla, CA, USA) to trigger Smok Baby Beast Brother TFV8 sub-ohm tanks. Vapor delivery was automated by computer and occurred in parallel across groups; thus, the experimenter was not blinded to Adolescent Drug treatment conditions. Tanks were equipped with V8 X-Baby M2 0.25 ohm coils. Vapor exposure chambers were attached to vacuum exhausts which pulled ambient air through intake valves at ∼1.5-2 L per minute. Twice daily (6 h inter-session interval) heroin (50 mg/mL in PG) or PG (control) vapor exposure sessions began on PND 36. Drug exposure conditions were derived from our prior work showing self-administration (Gutierrez et al. 2020), maximum nociception and virtually complete locomotor suppression (Gutierrez et al. 2021). Rats were treated in groups of six per chamber and exposure sessions consisted of six-second vapor deliveries separated by five-minute intervals. The house vacuum was turned on 30 seconds prior to vapor deliveries and closed immediately after. Total vapor clearance time was ∼5 minutes. Vapor began being cleared from the chamber 4.5 minutes after the last vapor delivery. Each vapor exposure chamber was dedicated to a single treatment group and was cleaned after each session using an alcohol solution. Vapor exposure sessions continued for total of 10 days (20 sessions; PND 36 -PND 45; **Fig 1**).

**Fig. 1.**
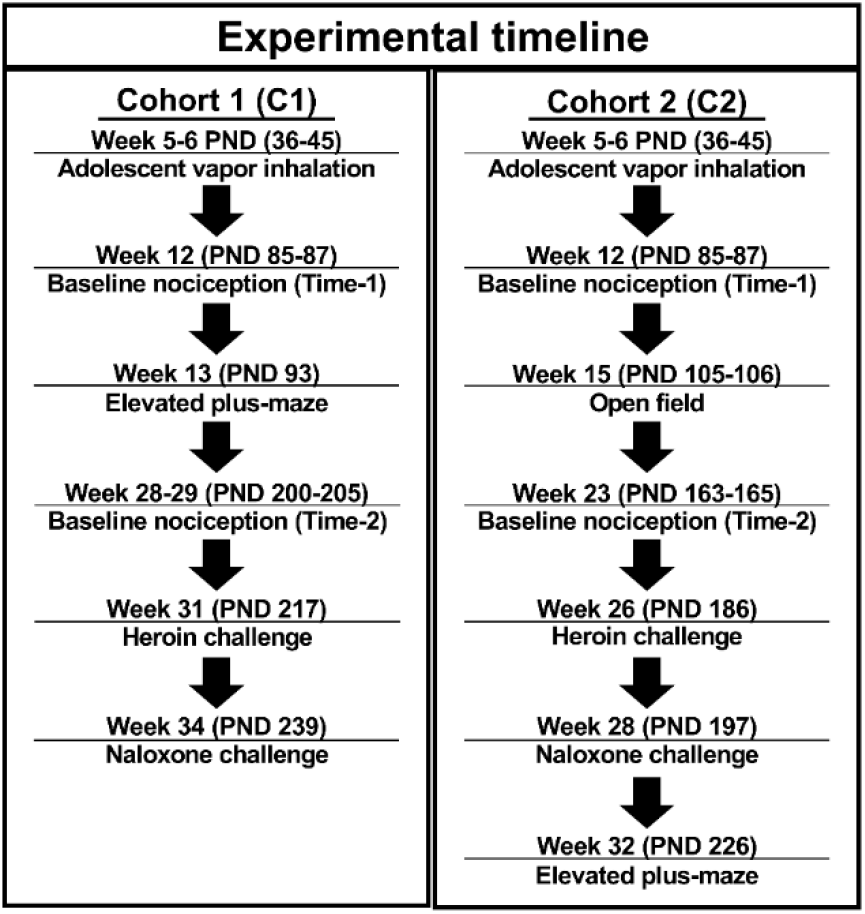
Experimental timeline for Cohort 1 (left panel; C1) and Cohort 2 (right panel; C2) rats.

### Thermal nociception

#### Nociception assays

A Branson Brainsonic CPXH Ultrasonic Bath (Danbury, CT) was used to perform tail-withdrawal tests. The bath was filled with water and the temperature was set and maintained at the target temperature (46°C, 48°C, 50°C, °r 52°C depending on the experiment). Target temperatures were selected based on our previous work (Gutierrez et al. 2021; Gutierrez et al. 2022; Gutierrez et al. 2020) and standard ranges of water temperatures for assessing nociception in rodents (Deuis et al. 2017). The water bath was stirred, and the temperature was verified using a second thermometer prior to each rat assessment. Water temperature was kept at the target temperature +/-0.10°C. A stopwatch was used to measure tail-withdrawal latency. One tail-withdrawal measurement was performed for each rat at each temperature assessed for a given time/age of measurement. A cutoff of 15 sec was used to avoid any potential for tissue damage.

#### Baseline nociception

The first set of nociception assays (Time-1) were performed at 12 weeks of age over two days (PND 85 and PND 87; **Fig 1**), with two temperatures assessed per test day (48°C and 52°C on one day and 46°C and 50°C on the other). Tail-withdrawal measurements for each test day were separated by 120 minutes, and a counterbalanced order was used for the two designated test temperatures. A range of water temperatures was used to ensure any between-groups differences were not obscured by ceiling or floor effects. A second set of tail-withdrawal measurements (Time-2) were performed at 23 weeks of age (PND 163 and PND 165; **Fig 1**) in C2 rats and 28-29 weeks of age (PND 200 and PND 205; **Fig 1**) in C1 rats. Assessments performed at a temperature of 46°C were carried out on test day one and those at 48°C and 52°C were carried out on test day 2. On this latter, the two measurements were separated by 120 minutes and a counterbalanced order was used for the two designated temperatures.

#### Heroin and naloxone challenge

The anti-nociceptive effects of acute heroin injection were assessed in C2 rats at 26 weeks of age (PND 186; **Fig 1**) and in C1 rats at 31 weeks of age (PND 217; **Fig 1**). Rats were administered heroin (0.5 mg/kg, s.c.) 30 minutes prior to the first tail-withdrawal measurement, and then injected with heroin (1.0 mg/kg, s.c.) 120 minutes after the first injection; tail-withdrawal latencies were once again measured after a 30-minute interval. Baseline tail-withdrawal latencies from the Time-2 re-assessments were used for baseline comparisons. Water temperature for both assessments was set to 52°C and a cutoff of 15 sec was used.

At 28 (PND 197; Cohort C2; **Fig 1**) or 34 (PND 239; Cohort C1; **Fig 1**) weeks of age, rats were administered naloxone (0.3 mg/kg, i.p.), followed by naloxone (1.0 mg/kg, i.p.) 120 minutes after the first injection. Tail-withdrawal testing was conducted at 48°C, 15 minutes after each antagonist injection. Baseline tail-withdrawal latencies for this experiment were measured immediately before the first naloxone injection. The 48°C temperature was selected to provide parametric range to observe any anticipated hyperalgesic effects of naloxone injection.

### Anxiety-like behavior

The effects of adolescent heroin inhalation on the expression of anxiety-like behavior were assessed using an Elevated plus-maze (EPM) test and by assessing behavior in an open field arena (see **Supplementary Information**). EPM testing was conducted on week 13 (PND 93; **Fig 1**) in C1 rats (N = 10) and week 32 (PND 226; **Fig 1**) in C2 rats (N = 11). One C2 female rat in the heroin Adolescent Drug group jumped off the EPM apparatus during testing and was not included in the analysis. Test sessions were conducted in a dimly lit room and animals’ activity was recorded for 5 minutes. The apparatus had two opposing open arms and two opposing closed arms perpendicular to the open arms. Arms measured 50 cm in length and 10 cm in width. The walls of the closed arms were 40 cm in height. The apparatus had four legs, each positioned towards the distal portion of an arm, which suspended it 50 cm above the floor. Behavior was recorded and measured using AnyMaze software (Stoelting, Wood Dale, IL). Open arm measures were considered separately from Center measures and were therefore analyzed as independent zone measures. The apparatus was cleaned with an ethanol solution prior to the testing of each subject.

### Data and statistical analysis

Data collected from cohort 1 and 2 animals during baseline nociception measurements, EPM testing, and from the heroin and naloxone challenges were combined for analysis. Data were analyzed by Analysis of Variance (ANOVA) with repeated measures factors for Time in weight data and open field data, for Temperature in tail-withdrawal experiments where multiple temperatures were assessed, and for Treatment Condition in the heroin and naloxone challenge experiments. Mixed-effects analysis was used when there were missing data. Significant effects were followed by post-hoc analysis using the Sidak (two-level factors) or Tukey (multi-level factors) methods. A criterion of p < 0.05 was used to infer significant effects. Analyses were performed using Prism 9 for Windows (GraphPad Software, Inc, San Diego, CA).

## Results

### Thermal nociception

Initial analysis of thermal nociception at 12 weeks of age (PND 85 and PND 87; **Fig. 2**) confirmed a main effect of Temperature (F (3, 126) = 222.4, p < 0.0001), an interaction of Sex with Adolescent Drug (F (1, 42) = 8.639, p < 0.01), and a three-way interaction of Temperature, Sex, and Adolescent Drug (F (3, 126) = 3.197, p < 0.05). The post-hoc analysis confirmed significant differences between all temperatures examined. Two-way ANOVAs were then performed at each level of Sex and Temperature to further parse effects. Within female groups, a significant effect of Adolescent Drug (F (1, 22) = 12.08, p < 0.01; **Fig. 2A**) was confirmed; adolescent heroin treatment increased withdrawal latencies. A significant effect of Temperature (F (3, 66) = 104.5, p < 0.0001; **Fig. 2A**) was also confirmed and the post-hoc analysis confirmed significant differences between all temperatures. Within male groups, a significant effect of Temperature (F (3, 60) = 121.7, p < 0.0001; **Fig. 2B**) was confirmed and the post-hoc analysis further confirmed significant differences between all temperatures examined. No main effect of Adolescent Drug was confirmed for the male groups (**Fig. 2B)**. When Adolescent Drug and Sex were considered at each level of Temperature, significant interactions of Adolescent Drug with Sex were confirmed at 46°C (F (1, 42) = 8.474, p < 0.01; **Fig. 2C**) and at 50°C (F (1, 42) = 5.164, p < 0.05; **Fig. 2D**). Post-hoc analyses confirmed a difference between PG- and heroin-treated females at 46° but not at 50°C. The 15 second cutoff duration for warm-water tail submersion during testing was only reached by 3 PG female, 6 PG male, 8 heroin female and 2 heroin male rats in the 46°C condition.

**Fig. 2.**
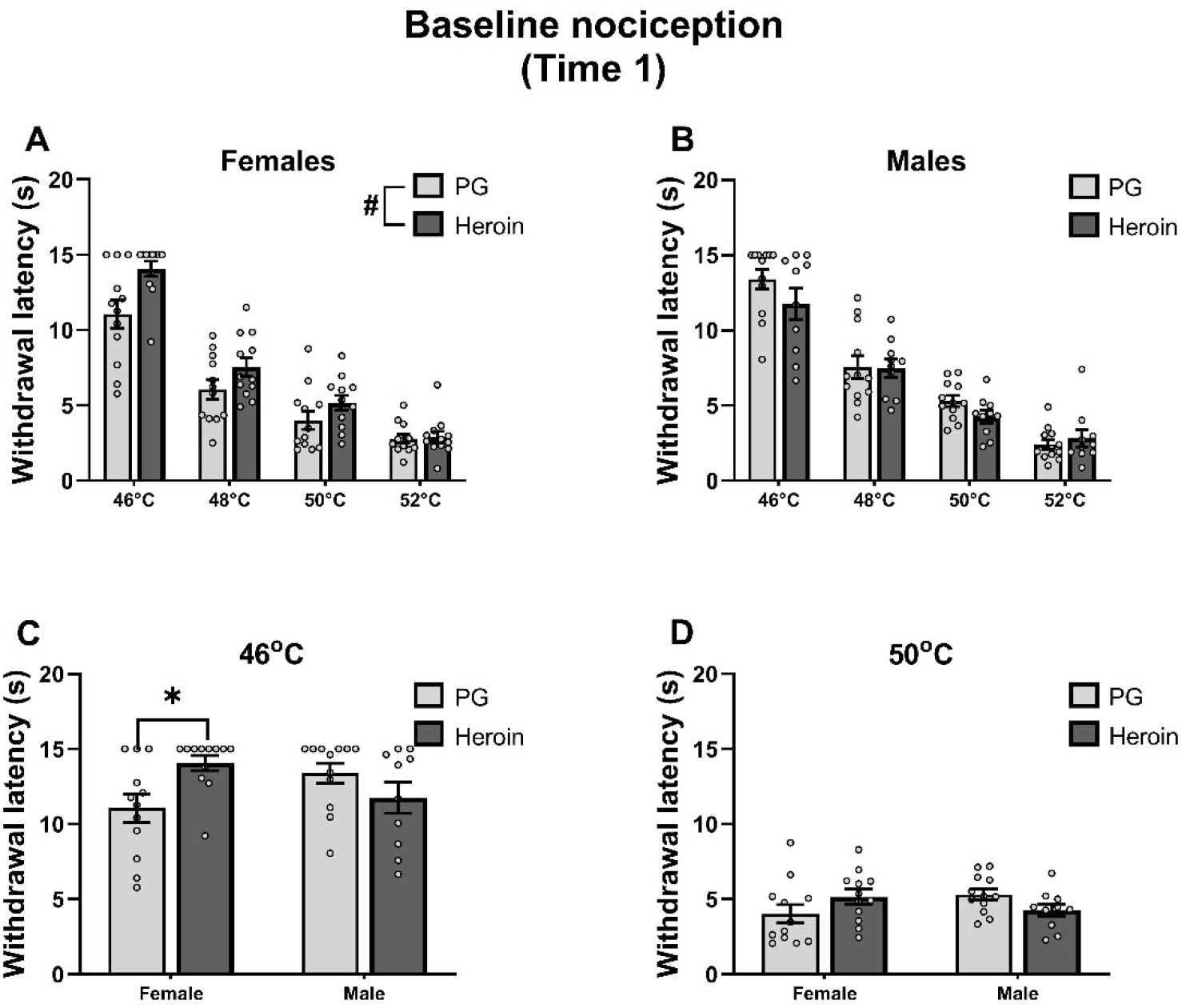
Mean (±SEM) baseline tail withdrawal latencies for female (N=24; 12 PG, 12 Heroin) and male (N=22; 12 PG, 10 Heroin) rats at 12 weeks of age (PND 85-87). Latencies for female (A) and male (B) rats, treated with PG or Heroin during adolescence, across four temperatures (46°C, 48°C, 50°C, and 52°C). Withdrawal latencies for female and male animals, treated with PG or heroin during adolescence, at 46°C (C) and 50°C (D). A significant difference between adolescent drug groups (within sex) collapsed across temperature with #; a difference between adolescent drug groups (within sex) for a given temperature with *.

The three-way analysis of baseline nociception at 23 weeks (PND 163-165; Cohort C2) and ∼29 weeks (PND 200-205; Cohort C1) of age (Time-2) confirmed significant effects of Temperature (F (2, 84) = 126.9, p < 0.0001; **Fig. 3A**) and Adolescent Drug (F (1, 42) = 6.757, p < 0.05; **Fig. 3A**) but not of Sex (**Fig. 3B and 3C**); no significant interactions of factors were confirmed. That is, heroin-treated animals exhibited significantly shorter latencies compared with PG-treated animals, collapsed across Temperature or Sex. For Temperature, latencies at 52°C were significantly lower compared with those at 46°C or 48°C and latencies at 48°C were significantly lower compared with those at 46°C.

**Fig. 3.**
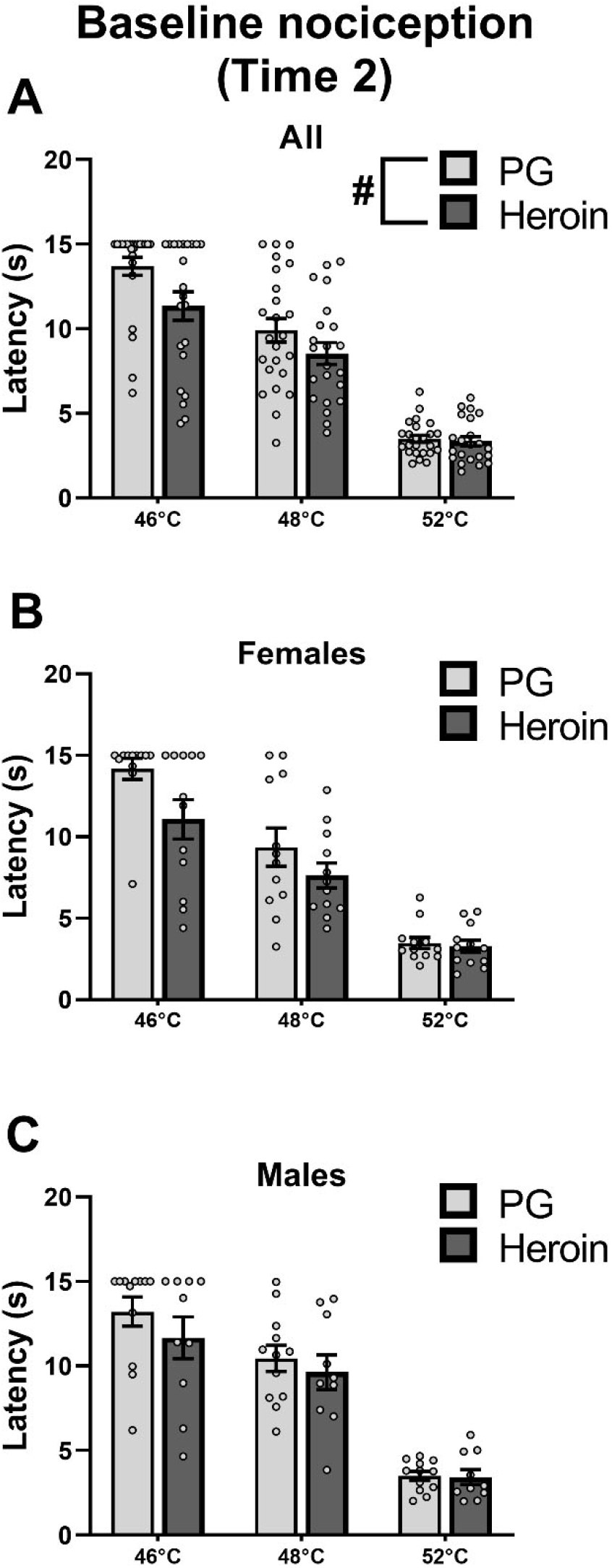
Mean (±SEM) baseline tail withdrawal latencies for female (N=24; 12 PG, 12 Heroin) and male (N=22; 12 PG, 10 Heroin) rats at the Time-2 (23/29 weeks) measurement. Withdrawal latencies collapsed across sex (A) and separated into female (B) and male (C) groups. A significant difference between groups (collapsed across sex and temperature) is indicated with #.

### Heroin and naloxone challenge

Injection of heroin (0.5, 1.0 mg/kg, s.c.) slowed tail withdrawal latencies in the nociception test. The three-way ANOVA confirmed significant effects of Treatment Condition (F (2, 84) = 191.4, p < 0.0001), Sex (F (1, 42) = 6.719, p < 0.05), and Adolescent Drug (F (1, 42) = 16.07, p < 0.001) on withdrawal latency; adolescent heroin treatment significantly decreased withdrawal latencies (**Fig. 4A**), female rats exhibited significantly shorter latencies compared with male rats (**Fig. 4B**), and all Treatment Conditions significantly differed from each other. Additionally, Adolescent Drug interacted significantly with Treatment Condition (F (2, 84) = 6.535, p < 0.01; **Fig. 4**).

**Fig. 4.**
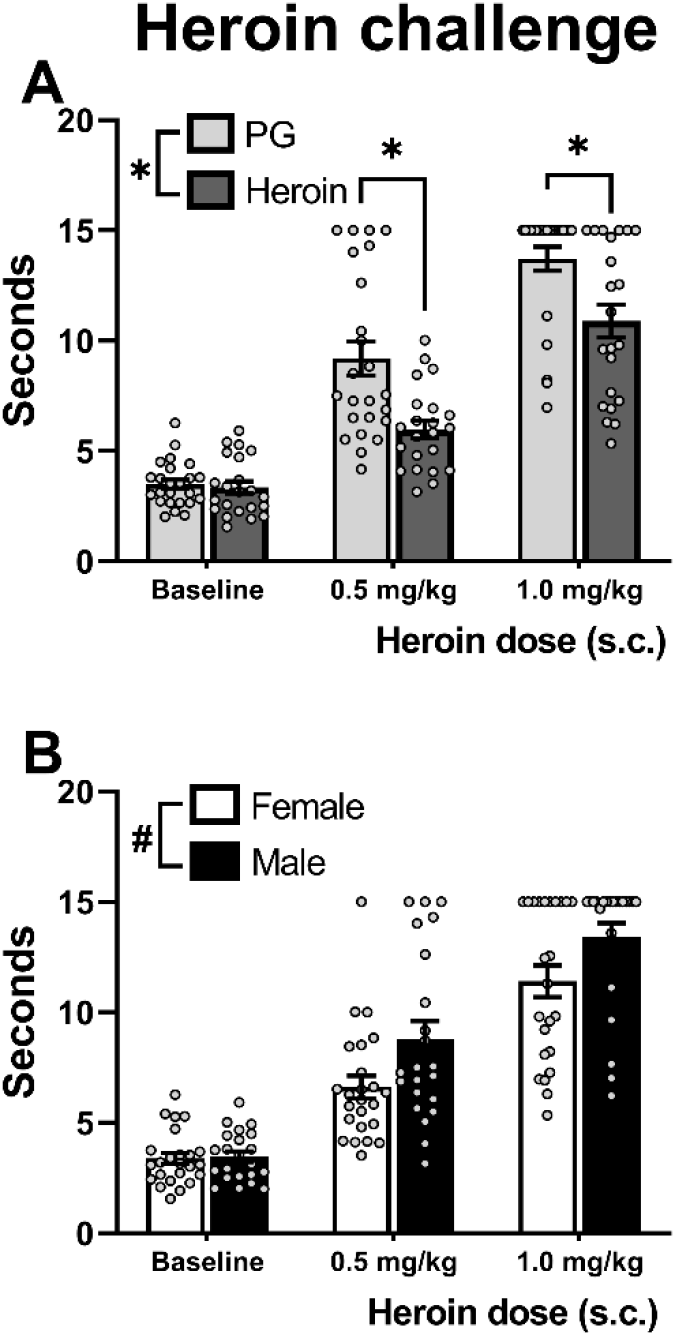
Mean (±SEM) tail withdrawal latencies at 52°C for female (N=24; 12 PG, 12 Heroin) and male (N=22; 12 PG, 10 Heroin) rats under baseline or Heroin challenge (0.5, 1.0 mg/kg, s.c.) conditions collapsed across Sex (A) and Adolescent Drug (B). A significant difference between Adolescent Drug groups (within treatment condition) is indicated with *; a difference between sexes with #.

The interaction was followed up by performing a two-way ANOVA collapsed across Sex. Significant effects of Adolescent Drug (F (1, 44) = 14.74, p < 0.001), Treatment Condition (F (2, 88) = 179.8, p < 0.0001), and the interaction of Adolescent Drug with Treatment Condition (F (2, 88) = 6.360, p < 0.01) were again confirmed. The post-hoc analysis confirmed significantly slower tail-withdrawal latencies for animals treated with heroin during adolescence, compared with those treated with PG, following the 0.5 mg/kg and the 1.0 mg/kg doses (**Fig. 4A**).

A three-way analysis of withdrawal latencies (48°C water) following naloxone injections confirmed a significant effect of Treatment Condition (F (2, 84) = 7.168, p < 0.01; **Fig. 5**). The post-hoc analysis confirmed a significant decrease in latencies following treatment with 0.3 or 1.0 mg/kg naloxone compared with baseline. Additionally, there was a significant interaction of Treatment Condition with Sex (F (2, 84) = 3.423, p < 0.05). A two-way follow up analysis was then performed by collapsing across Adolescent Drug. Significant effects of Treatment Condition (F (2, 88) = 7.389, p < 0.01) and of the interaction of Treatment Condition with Sex (F (2, 88) = 3.552, p < 0.05) were again confirmed. Post-hoc analysis of the interaction confirmed a significant difference between female and male rats at baseline which appeared to be driven by the C1 PG-exposed female group exhibiting faster withdrawal latencies than any other group at baseline (**Supplementary Information**; **Fig. S2**). Latencies were also faster than those recorded for this group in the original evaluation at 48°C (**Fig. 3B**), unlike other groups, potentially compromising any detection of hyperalgesic effects of naloxone. The reason for this is unknown. Additionally, the post-hoc analysis of the interaction of Treatment Condition with Sex confirmed a significant difference between each naloxone treatment condition and baseline in the male rats, but not the female rats. A full depiction of tail-withdrawal latencies by Group and Cohort can be found in the **Supplementary Information** (**Fig. S2**); importantly, inclusion/exclusion of the C1 female groups has no differential impact on the statistical, and therefore interpretive, outcome for this experiment (see **Supplementary Information**).

**Fig. 5.**
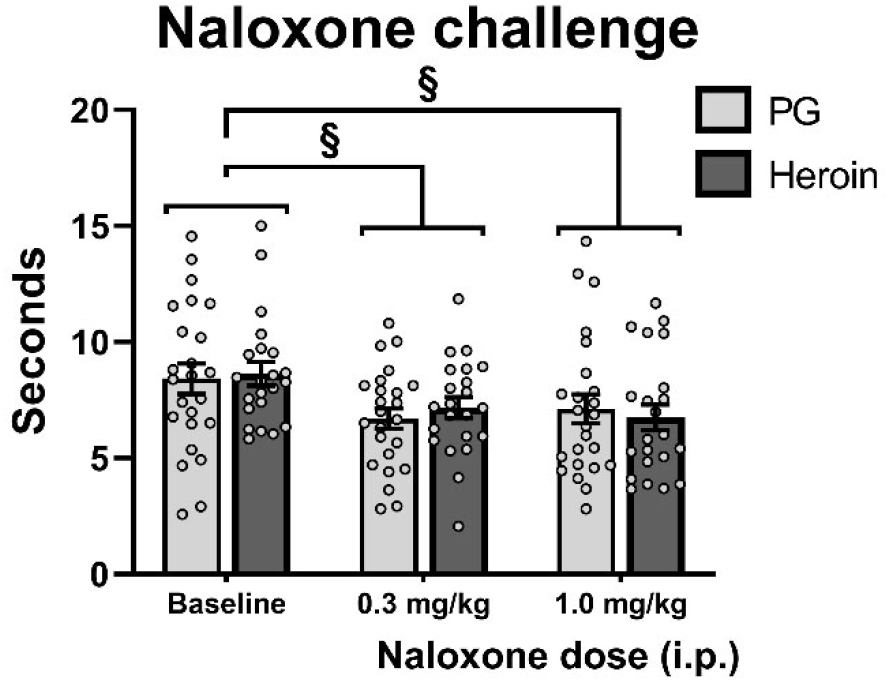
Mean (±SEM) tail-withdrawal latencies at 48°C for PG (N=24; 12 female, 12 male) and Heroin (N=22; 12 female, 10 male) groups under baseline or naloxone challenge (0.3, 1.0 mg/kg, i.p.) conditions collapsed across Sex. A significant difference between treatment conditions is indicated with §.

### Anxiety-like behavior

Analysis of time spent in the open arms of the elevated plus-maze confirmed significant effects of Adolescent Drug (F (1, 41) = 5.632, p < 0.05; **Fig. 6A**) and of Sex (F (1, 41) = 5.442, p < 0.05; **Fig. 6A**). Adolescent heroin treatment significantly reduced open arm time, and males spent significantly less time in the open arms compared with females. Significant effects of Sex, but not Adolescent Drug, were also confirmed for mean speed ((F (1, 41) = 18.21, p < 0.0001; **Fig. 6B**), total distance traveled ((F (1, 41) = 18.52, p < 0.0001; **Fig. 6C**), overall immobility ((F (1,41) = 49.56, p < 0.0001; **Fig. 6D**), and total number of arm entries (F (1, 41) = 5.489, p < 0.05; not shown). Female rats were significantly faster, traveled more distance, had a higher number of total arm entries (mean entries: 31.93 female, 26.15 male), and spent significantly less time immobile compared with male rats. No differences in Open arm entries or Time spent in the Center zone were confirmed Separate three-way analyses including Cohort (and therefore age) as a factor were performed for Time spent in the Open Arm zones, average Speed, total Distance traveled, number of Entries into the Open Arm zones, overall Immobility, and Time spent in the Center zone to assess the impact of conducting the elevated plus-maze test at different ages in the two cohorts (see **Supplementary Information**). A significant effect of Adolescent Drug was only confirmed for Time spent in the Open arms (see **Supplementary Information**; **Fig. S3**). Testing the Cohorts at different ages did not have an impact on differences associated with Adolescent Drug treatment in the Open arms, i.e., there was no main effect of Age confirmed for Time spent in the Open Arms, nor any interaction of Age with Adolescent Drug or with Sex. However, there was a significant interaction of Age with Sex (F (1, 37) = 19.23, p < 0.0001).

**Fig. 6.**
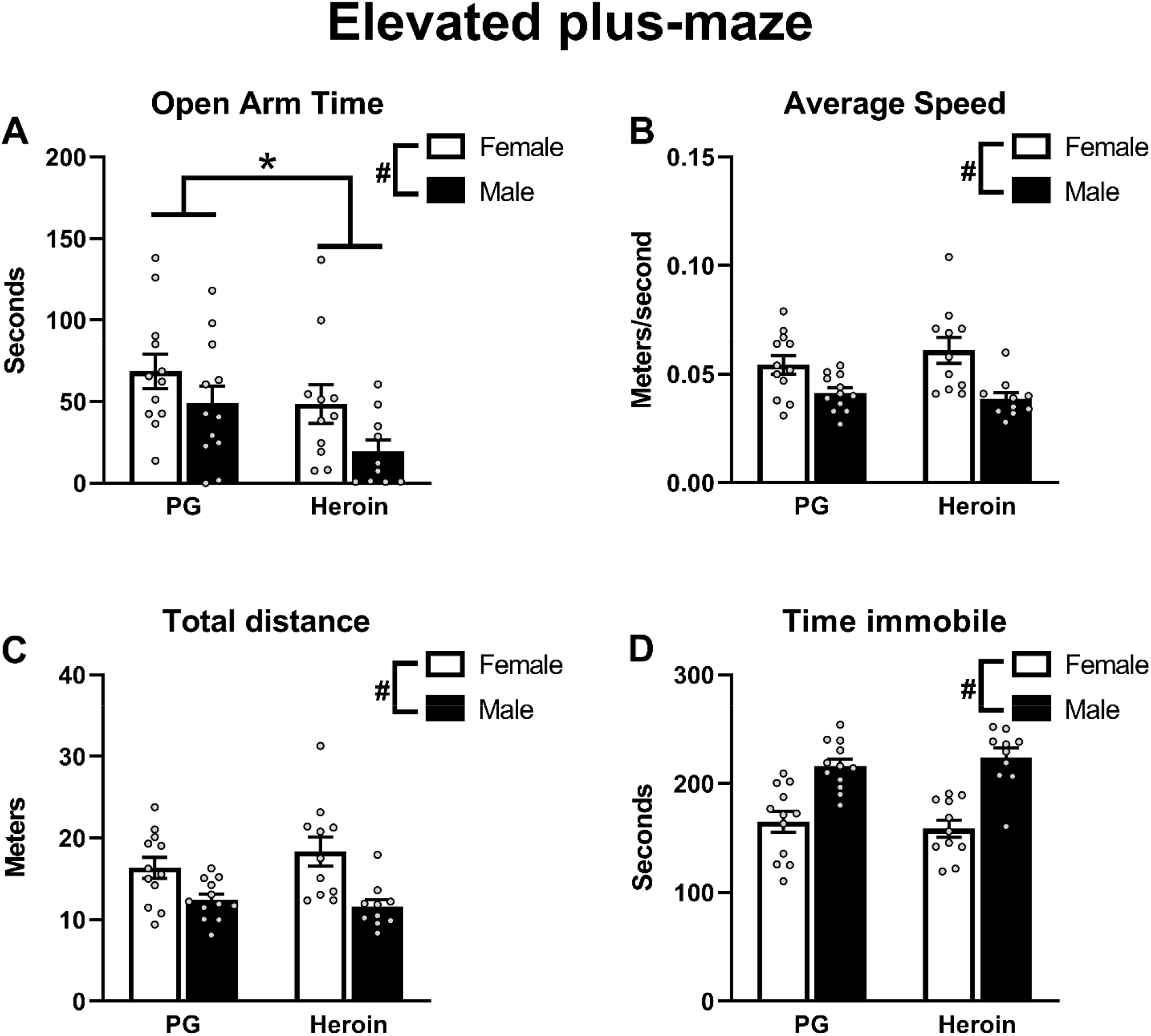
Mean (±SEM) A) time spent in the open arms, B) speed of movement, C) distance traveled and D) time spent immobile in the Elevated Plus-Maze test for female (N=23; 12 PG, 11 Heroin) and male (N=22; 12 PG, 10 Heroin) rats in the elevated plus-maze. A significant difference between adolescent drug groups with *; a difference between sexes with #.

The post-hoc analysis confirmed that female rats tested at 32 weeks of age (PND 226) spent significantly *more* time in the Open arms compared with 13-week (PND 93) old female rats whereas 32-week old male rats spent significantly *less* time in the Open arms compared with 13-week old male rats (**Fig. S3**). Furthermore, 32-week old female rats spent significantly more time in the Open arms compared with male rats at 32 weeks of age, but there were no significant differences between sexes at the 13-week assessment.

Open field behavior data are reported in the **Supplementary Information**.

## Discussion

Repeated exposure to heroin vapor during adolescence produced behavioral and physiological changes that lasted into adulthood in both female and male rats, in this study. The data confirmed adolescent heroin inhalation decreased sensitivity to thermal nociception in female rats upon reaching early adulthood, but increased sensitivity in both female and male rats when re-assessed at 23 and 29 weeks of age. This hyperalgesia was further confirmed by a reduction in anti-nociceptive efficacy of acute heroin injections in the groups exposed to heroin vapor. Expression of increased anxiety-like behavior was also observed in the elevated plus-maze test in female and male rats exposed to heroin. This is the first study to report long-term hyperalgesia, decreased opioid anti-nociceptive efficacy, or anxiety-like behavior in adult animals exposed to opioids as adolescents where inhalation was used as the experimental route of drug administration. In humans, inhaled heroin produces more positively rated subjective effects compared with those produced by intravenous administration, despite lower C_MAX_ heroin levels (Rook et al. 2006). While this may suggest that heroin inhalation is protective compared with intravenous administration, there are risks specific to route such as toxic leukoencephalopathy associated with “chasing the dragon” (Alambyan et al. 2018; Cheng et al. 2019). There are also pharmacokinetic differences between routes, such as an increase in the heroin metabolite morphine-6-glucuronide when heroin is inhaled (Rook et al. 2006) and, of course, rapid-offset of subjective drug effects can result in frequent re-dosing and binging behavior. The contributions of such differences to the lasting consequences of opioid exposure are presently not well understood, therefore the findings reported here provide important information about how the inhalation method of exposure can potentially impact outcome.

Lasting, potentially life-long, increases in anxiety and a need for higher opioid doses for effective analgesia caused by adolescent opioid inhalation may be risk factors for developing later life opioid use disorders (Martins et al. 2012; Rogers et al. 2019).

The present study found sex-specific effects of adolescent heroin exposure on baseline nociception at 12-weeks, but this shifted qualitatively by the time the second basal measurement was performed. An initial *hypo*algesia in the heroin-exposed female group changed to the *hyper*algesia that was present in the male group at both time points. Sex-specific analgesic effects of opioids have been documented in pre-clinical studies. For example, morphine produces greater acute anti-nociceptive effects and greater tolerance following chronic treatment in male rats (Craft et al. 1999). Additionally, continuous opioid agonist infusion produces hyperalgesia faster and for a longer duration in female mice, but at higher doses persists equally and is prevented only in males by an N-Methyl-d-aspartate (NMDA) antagonist (Juni et al. 2008). The reasons for this interesting finding within the present study are not known and will require future studies to better understand the cause and implications of the qualitative shift in nociceptive sensitivity observed in heroin-exposed female rats from 12 weeks to 23/29 weeks of age. It may be the case, for example, that male rats would also exhibit hypoalgesia if assessed at a timepoint closer to the adolescent heroin exposure.

Opioid tolerance and opioid-induced hypersensitivity to noxious stimuli are commonly reported adverse effects of repeated opioid treatment (Allouche et al. 2014; Célèrier et al. 2001; Colvin et al. 2019; Costa et al. 2020; Doyle et al. 2020; Gardell et al. 2006; Vanderah et al. 2001). The development of tolerance to opioids can lead to dose escalation in patients in clinical settings (Han et al. 2013) and in illicit opioid users (Cowan et al. 2001). Opioid dose escalation for the management of pain can result in opioid-induced hyperalgesia, thereby worsening clinical outcomes (Mercadante et al. 2003), and opioid discontinuation in dependent users can result in decreased pain thresholds, which can last for months (Carcoba et al. 2011). Increased pain sensitivity is commonly reported among abstinent users and former opioid abusers in opioid maintenance programs, and is associated with heightened opioid craving (Compton et al. 2001; Ren et al. 2009; Tsui et al. 2016), which increases the risk of relapse. In the present study, animals exposed to heroin by vapor inhalation as adolescents displayed increased thermal sensitivity when assessed starting at 23 weeks of age, and a reduction of anti-nociceptive efficacy of acute heroin injections. It is not clear, however, if this was due to an apparent tolerance, e.g. increased NMDA receptor mediated pronociceptive activity can mask the anti-nociceptive effects of the injected heroin (Laulin et al. 1999), or if this was due to persistent changes in opioid receptor function (Allouche et al. 2014). Interestingly, when injected with naloxone tail-withdrawal latencies were generally decreased, but there was no exacerbated response observed in the heroin-treated rats. Heightened endogenous mu-opioid anti-nociceptive activity, in response to heightened pro-nociception (Célèrier et al. 2001), may therefore not necessarily be responsible for the behavioral observations. Determining the mechanisms responsible for the persistent nociceptive effects observed here is outside the scope of this study, however, and will require future studies to delineate the neurobiological substrates underlying the reported behaviors.

The relationship between substance abuse and anxiety disorders is a complex and cyclical one. The two often co-exist (George et al. 1990; Kushner 2000), anxiety disorders during adolescence increase the later risk of substance abuse disorders (Bushnell et al. 2019; Lopez et al. 2005), and adolescent substance use increases the risk of anxiety disorders later in life (Brook et al. 1998; Moylan et al. 2013). Here we examined the effects of adolescent heroin vapor exposure on the expression of an anxiety-like behavioral phenotype in adult male and female rats, using an EPM approach. In contrast to a previous study reporting no alteration of anxiety-like behavior in an EPM test in adult rats repeatedly treated with morphine, i.p., as adolescents (Schwarz and Bilbo 2013), the present study found that adolescent heroin vapor exposure decreased time spent in the open arms of the test apparatus; a decrease in time spent by an animal in the open areas of the EPM (or similar behavioral tests such as Elevated Zero Maze) is widely interpreted as an increase in anxiety-like behavior in rodents (Lister 1987; Pellow et al. 1985; Sprowles et al. 2016). Importantly, the expression of anxiety-like behavior observed in this study extended from early to middle adulthood, long after the initial heroin exposure during adolescence. The impact of adolescent heroin was consistent across sex, despite the fact that male rats exhibited significantly higher overall immobility, lower average speed, and traveled less distance compared with female rats.

The EPM and open field test are both commonly used to assess anxiety-like behavior related to opioid drug treatment in rodents. For example, morphine withdrawal generally reduces open arm time and entries in the EPM (Bhattacharya et al. 1995; Schulteis 1998; Zhang and Schulteis 2008), although open arm time and entries in the EPM have been reported in mice following spontaneous, precipitated, and protracted morphine withdrawal (Bravo et al. 2020; Buckman et al. 2009; Hodgson et al. 2008; Liptak et al. 2012). Similarly, morphine withdrawal reduces center activity in the open field test (Raghav et al. 2021). Interestingly, while some studies that have used the EPM and open field tests in conjunction have confirmed effects by both tests, in other studies the effects that have been identified by one test have not always been confirmed by the other. For example, anxiolytic and anxiogenic effects were reported in both tests in rats two hours and two days after the last of a series of daily morphine injections, respectively (Avila et al. 2008; Wang et al. 2017). In contrast, treatment with κ-opioid receptor (KOR) antagonists increased open arm measures in the elevated plus maze but had no effect on open field activity (Knoll et al. 2007), and a small dose of a KOR agonist produced anxiolytic effects in the EPM without impacting open field behavior (Privette and Terrian 1995) in rats. In the current study, we confirmed effects of adolescent heroin exposure on measures of anxiety-like behavior in the EPM but not in the open field. The reasons for this are unclear but could potentially involve differences in test sensitivity to specific components of anxiety. Indeed, there is evidence that suggests that the two may test different aspects of anxiety (Radhakrishnan and Gulia 2018; Sudakov et al. 2013), and one meta-analysis reported compelling evidence that sensitivity of the open field test as a test of anxiety may depend on specific pharmacology, and therefore on the specific model of anxiety (Prut and Belzung 2003). It is possible that the EPM may have been a more sensitive test to the specific behavioral effects produced by our inhalation model. Furthermore, the lack of treatment effects related to activity in the open field may actually reinforce our findings in the EPM by suggesting that anxiety-like behavior observed in heroin-treated animals in the EPM is not likely driven by alterations in overall activity levels. Additionally, while no differences were confirmed in open field behavior between adolescent vapor treatment groups, sex differences in overall activity *were* observed in the open field test (**Fig. S4A**), as were differences in activity specific to the center region of the apparatus (**Fig. S4B**). The fact that these findings are in agreement with the well-documented sex-dependent differences in activity in rodents (Belviranli et al. 2012; Simpson and Kelly 2012) serves to further validate the EPM and open field methodology used in this study.

There are a number of considerations regarding the novel vapor exposure paradigm that deserve discussion. One such consideration is the degree of drug uptake, given that there are a number of factors such as activity level during dosing, respiration rate, and location of an animal within the inhalation chamber that could have a potential impact. Nevertheless, our previous published behavioral data comparing injected with inhaled heroin, using the same parameters, show similar degree of inter-subject variability and magnitudes of acute behavioral effects between the two methods (Gutierrez et al. 2021). Therefore, blood heroin levels after the vapor exposure described here is most likely commensurate with those achieved in the more traditional parenteral routes of administration. Secondly, it is possible the PG vehicle may itself produce effects. Two prior reports found anxiolytic effects of PG administration (Da Silva and Elisabetsky 2001; Lin et al. 1998), however, those studies report acute effects in mice that received PG *by injection*. At present, no data exist that suggest long-term effects of PG vapor inhalation on anxiety-like behavior. Lastly, it should be noted that animals were allowed only six days of habituation from arrival at the vivarium to the start of vapor inhalation sessions. It is not known what effect this may or may not have had on dosing.

In summary, the findings of this study show that lasting behavioral and physiological changes are produced by repeated heroin vapor exposure during adolescence in rats. This included a demonstration that long-term changes in nociception persist deep into adulthood, a phenomenon that is not well-described in the literature. Additionally, these results suggest that sex, at least during early adulthood, may play a determinative role in the manner by which the nociceptive effects of repeated heroin exposure are expressed. The present study also shows that long-lasting anxiety-like effects are produced by repeated adolescent heroin exposure. Finally, these data show for the first time that heroin inhalation is capable of inducing these persisting, possibly life-long, alterations in anxiety and nociception. This further validates use of EDDS technology for the study of the effects of inhaled opioid drugs in rodents. We conclude that adolescent heroin inhalation may increase risk for later life development of opioid use disorder via reduced sensitivity to opioid analgesia and increased anxiety.

## Acknowledgements

This work was supported by USPHS grants (R01 DA035281, R01 DA042211 and T32 AA007456), a UCSD Chancellor’s Post-doctoral Fellowship (AG) and the Tobacco Related-Disease Research Program (TRDRP; T31IP1832). The National Institutes of Health, NIDA and NIAAA had no influence on the design, conduct, analysis and interpretation of the studies, nor on the decision to publish the findings. The authors declare no conflicts of interest.

