## Supplementary Information for "The long-term effects of repeated heroin vapor inhalation during adolescence on measures of nociception and anxiety-like behavior in adult Wistar rats"

### SUPPLEMENTAL METHODS

#### Open field activity

Open field activity was evaluated in Cohort 2 (C2) female (N = 6) and male (N = 6) rats at 15 weeks of age. Animals were recorded by video camera (Logitech C270) for a total of 60 minutes, however only the first 10 minutes were evaluated. Tests were conducted in plastic chambers with dimensions of 80 cm (L) X 44 cm (W) X 33 cm (H). Analysis of videographic recordings were conducted using ANY-maze software (Stoelting, Wood Dale, IL). Behavioral tracking was configured to distinguish activity in the Center (43 cm X 11 cm) versus the Peripheral zones.

Open field activity was again evaluated in C2 rats following acute heroin (s.c.) injection from 18 to 19 weeks of age. Rats were injected with saline or heroin (0.25, 0.5, or 1.0 mg/kg, s.c.) in a counter-balanced order. Rats were placed in the open field arena following injections and were recorded by video camera (Logitech C270) for a total of 60 minutes. Sessions were conducted up to twice per week, i.e., no more frequently than every 3-4 days. The data for this experiment were not reported here.

### SUPPLEMENTAL RESULTS

#### Weight

Rats were weighed weekly from 5 weeks of age to 39 weeks of age (**Fig. S1**). Weights for Cohort 1 (C1) animals for week 7 were recorded but were lost due to corruption of the storage device. A three-way analysis confirmed main effects of Time ($F(34, 1404) = 1079$, $p < 0.0001$) and Sex ($F(1, 42) = 545.6$, $p < 0.0001$) and an interaction of Time with Sex ($F(34, 1404) = 213.5$, $p < 0.0001$) but no effects of Adolescent Drug condition. The post-hoc analysis confirmed that male rats weighed significantly more compared with females at all time points except for the first recorded measurement at 5 weeks of age.

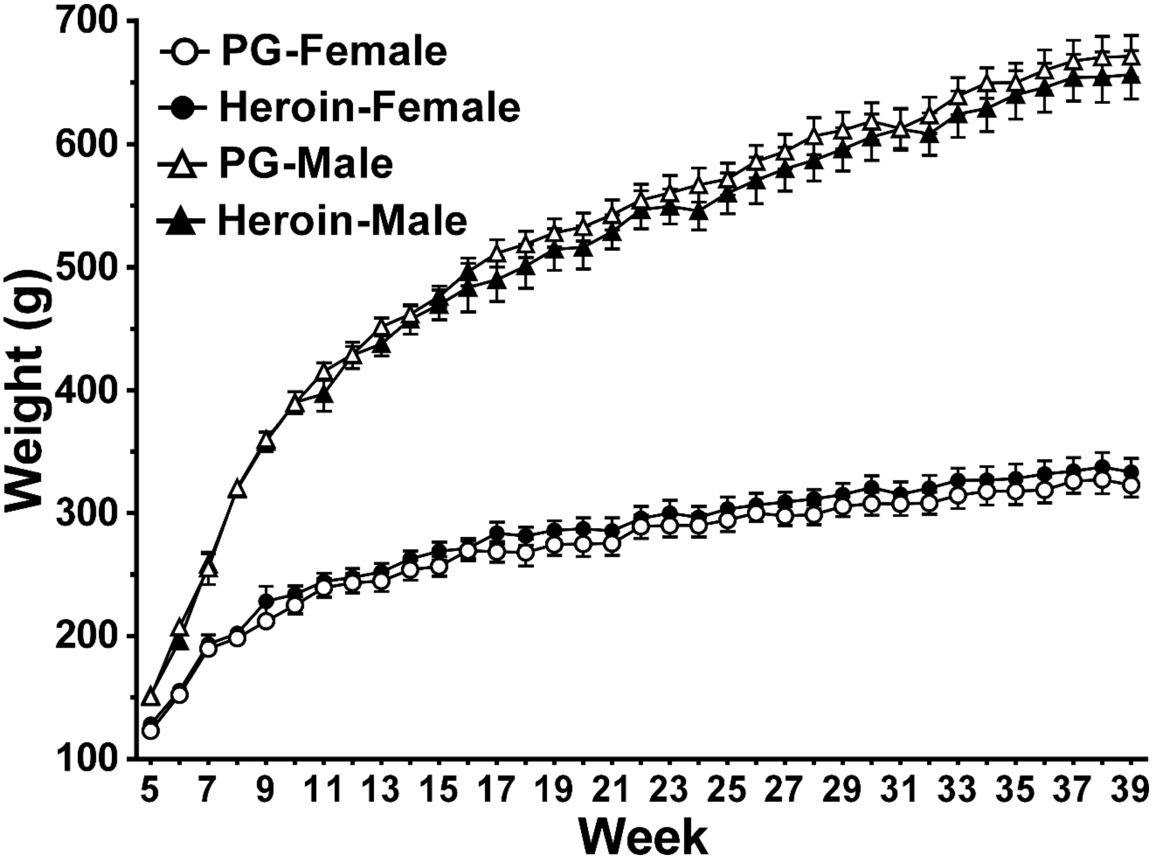


***Fig. S1*** *Mean (±SEM; N=12 per group) bodyweight for male and female rats (Weeks 5 – 39) exposed to repeated vapor from the propylene glycol (PG) vehicle or Heroin (50 mg/mL) during adolescence.*

**Naloxone challenge**

Separate three-way analysis were performed on tail-withdrawal latency data for each cohort in addition to the analysis performed on data from both cohorts combined (reported here and in the main **Results**) for the naloxone challenge experiments. Additionally, a three-way analysis was performed on tail-withdrawal latency data from both cohorts combined but with females from C1 excluded from the analysis. The three-way analysis for the Cohort 1 data confirmed only an interaction of Treatment Condition with Sex ($F(2, 36) = 4.115$, $p < 0.05$; **Fig. S2-C**). The post-hoc confirmed that the 0.3 mg/kg and the 1.0 mg/kg naloxone treatment conditions were each significantly decreased compared with the baseline condition in the males (**Fig. S2B**), but not the females (**Fig. S2A**). The three-way analysis of the Cohort 2 data confirmed a main effect of Treatment Condition (F (2, 40) = 16.31, p < 0.0001; **Fig. S2D-F**). The post-hoc confirmed that tail-withdrawal latencies for the 0.3 mg/kg and the 1.0 mg/kg naloxone treatment conditions were each significantly decreased compared with the baseline condition but were not significantly different compared with each other (**Fig. S2F**). The three-way analysis for

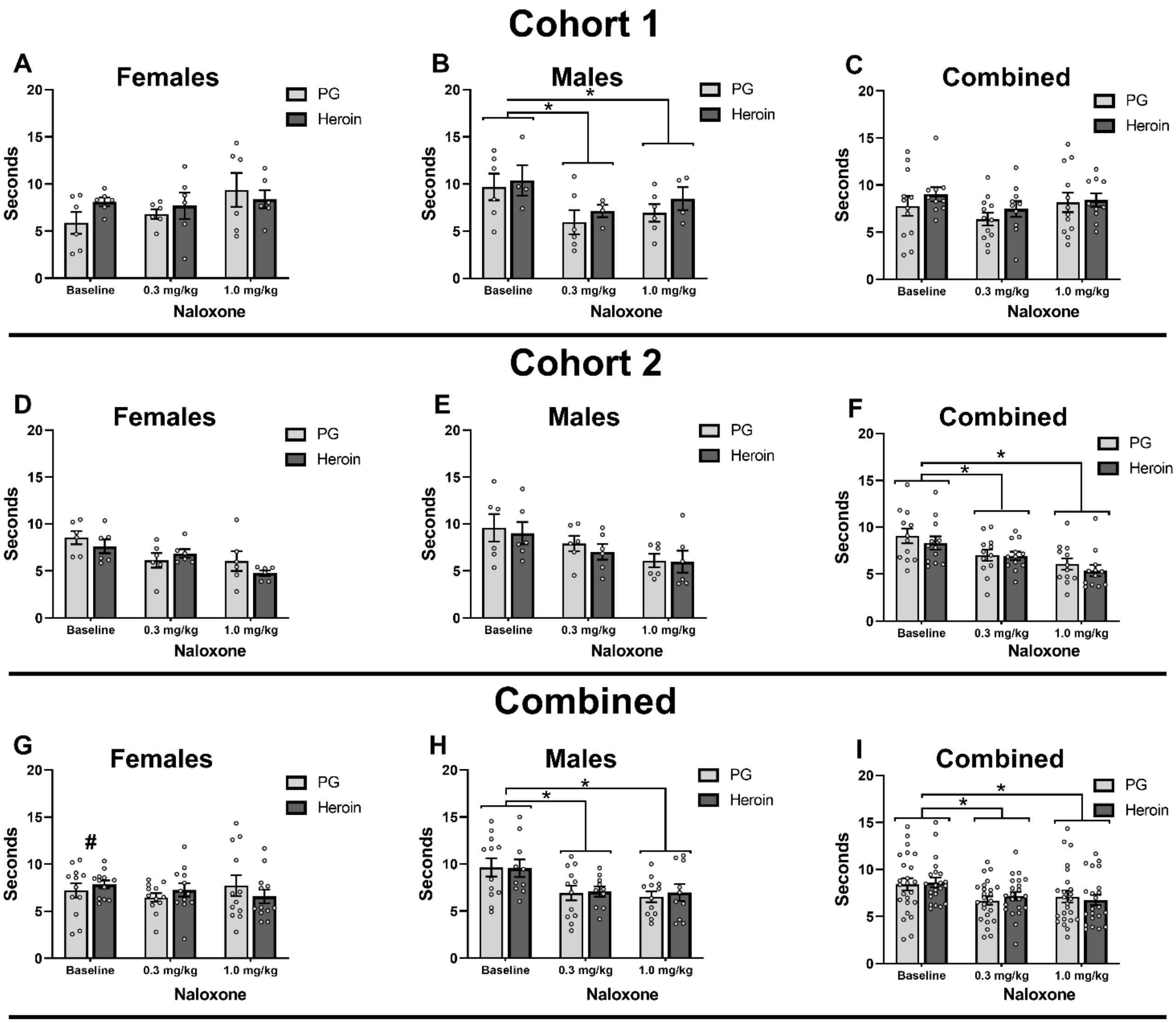


***Fig. S2*** *Mean (±SEM; Cohort 1: N=22, 12 female, 10 male; Cohort 2: N=24, 12 female, 12 male) tail-withdrawal latency for female and male rats at 48°C under baseline or naloxone challenge (0.3, 1.0 mg/kg, i.p.) conditions for the adolescent treatment groups. Withdrawal latencies for Cohort 1 females (A), males (B), and combined sexes (C). Withdrawal latencies for Cohort 2 females (D), males (E), and the combination (F). Withdrawal latencies for combined Cohort 1 and 2 females (G), males (H), and the combination of females and males from both cohorts (I). Significant differences between groups are indicated with *. Significant differences between sexes within Treatment Condition are indicated with #.*

data from the combined cohorts (**Fig. S2G-I**) confirmed a main effect of Treatment Condition ($F(2, 84) = 7.168$, $p < 0.01$; **Fig. S2I**) and an interaction of Treatment Condition with Sex ($F(2, 84) = 3.423$, $p < 0.05$). Follow-up post-hoc analysis of the main effect confirmed that the 0.3 mg/kg and the 1.0 mg/kg naloxone treatment conditions were each significantly decreased compared with the baseline condition (**Fig. S2I**). The follow-up analysis of the interaction, by collapsing across Adolescent Drug, once again confirmed significant effects of Treatment Condition ($F(2, 88) = 7.389$, $p < 0.01$) and of the interaction of Treatment Condition with Sex ($F(2, 88) = 3.552$, $p < 0.5$). The post-hoc analysis of the interaction confirmed that the 0.3 mg/kg and the 1.0 mg/kg naloxone treatment conditions were each significantly decreased compared with the baseline condition, but only in the males, and that baseline female latencies were significantly faster compared with male baseline latencies (**Fig. S2G, H**). The three-way analysis on the combined tail-withdrawal latency data with C1 females excluded from the analysis confirmed a main effect of Treatment Condition ($F(2, 60) = 17.01$, $p < 0.0001$). The post-hoc confirmed that the 0.3 mg/kg and the 1.0 mg/kg naloxone treatment conditions were each significantly decreased compared with the baseline condition.

**Elevated plus-maze**

Separate three-way analyses including Age as a factor were performed on Elevated plus-maze (EPM) data collected for Time spent in the Open Arm zones, average Speed, total Distance traveled, number of Entries into the Open Armzones, overall Immobility, and Time spent in the Center zone. Adolescent Drug effects were confirmed for Open arm behavior that were not influenced by factors of Sex or Age of testing. The analysis of Time spent in the Open arms confirmed effects of Adolescent Drug ($F(1, 37) = 5.975$, $p < 0.05$) and of Sex ($F(1, 37) = 6.878$, $p < 0.05$). Animals repeatedly exposed to heroin vapor as adolescents spent significantly less time in the Open arm zones compared with PG control animals, and female rats spent significantly more time in the Open arms compared with males. An interaction of Sex with Age ($F(1, 37) = 19.23$, $p < 0.0001$) was also confirmed (**Fig. S3**). The post-hoc analysis confirmed that female rats tested at 32 weeks of age spent significantly more Time in the

Open arm zones compared with females at 13 weeks of age while male rats at 32 weeks of age exhibited a significant decrease in Open arm Time compared with males at 13 weeks of age. Additionally, 32-week old female rats spent significantly more Time in the Open arms compared with 32-week old male rats. No differences were confirmed for Open arm Time between sexes at the 13-week assessment. Adolescent Drug did not interact with Sex or with Age in the analysis of Time spent in the Open arms. Analysis of average Speed confirmed effects of Age of testing ($F(1, 37) = 13.30$, $p < 0.001$) and of Sex ($F(1, 37) = 33.42$, $p < 0.0001$). Female rats were significantly faster compared with male rats, and rats tested at 32-weeks of age were significantly faster compared with those tested at 13 weeks of age. Age also interacted with Sex ($F(1, 37) = 18.61$, $p < 0.0001$). The post-hoc confirmed that female animals tested at 32 weeks of age were significantly faster than females tested at 13 weeks and males tested at both ages. No effects of Adolescent Drug were confirmed for average Speed. For distance traveled, the analysis confirmed effects of Age of testing ($F(1, 37) = 13.26$, $p < 0.001$) and of Sex ($F(1, 37) = 33.97$, $p < 0.0001$). Older rats traveled more distance compared with younger rats and female rats covered more distance compared with male rats. An interaction of Age with Sex ($F(1, 37) = 18.55$, $p < 0.0001$) was also confirmed. The post-hoc confirmed that, just as with Speed, female animals at 32 weeks of age traveled significantly more Distance compared with 13-week old female rats and with male rats at both ages. The analysis of Entries into Open arm zones confirmed an effect of Sex ($F(1,37) = 12.33$, $p < 0.01$). Female rats made

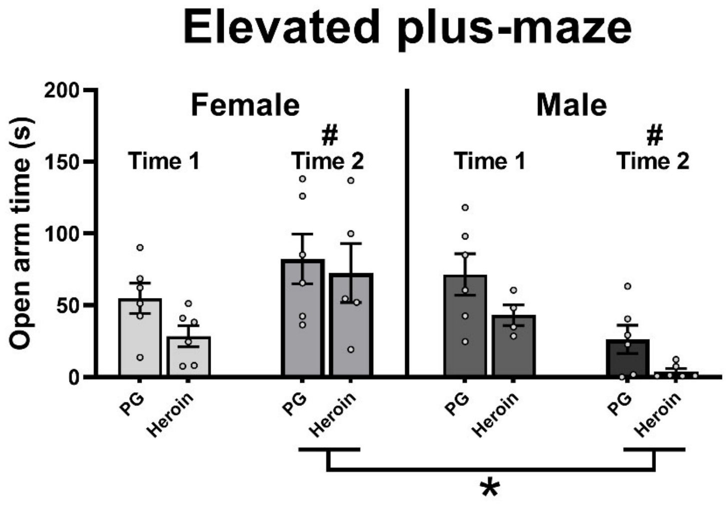


***Fig. S3*** *Mean (±SEM; Time 1: N=22, 12 female, 10 male; Time 2: N=23, 11 female, 12 male**)** Time spent in the Open arms of the elevated plus-maze. A significant difference compared with Time 1, within sex, is depicted with #; a difference between sexes at Time 2 with *.*

significantly more Entries into the Open arms compared with male rats. An interaction of Age with Sex ($F (1,37) = 15.24$, $p < 0.001$) was also confirmed. The post-hoc confirmed that 32-week old female rats entered into the Open arm zones significantly more compared with 13-week old female rats and with males at both ages tested. Analysis of overall Immobility confirmed effects of Age ($F (1, 37) = 9.713$, $p < 0.01$) and Sex ($F (1, 37) = 56.98$, $p < 0.0001$). Older animals were significantly more immobile compared with youngers animals, and male rats exhibited significantly more immobility compared with female rats. No effects of Adolescent Drug were confirmed for Immobility. There were no differences confirmed for Time spent in the Center zone

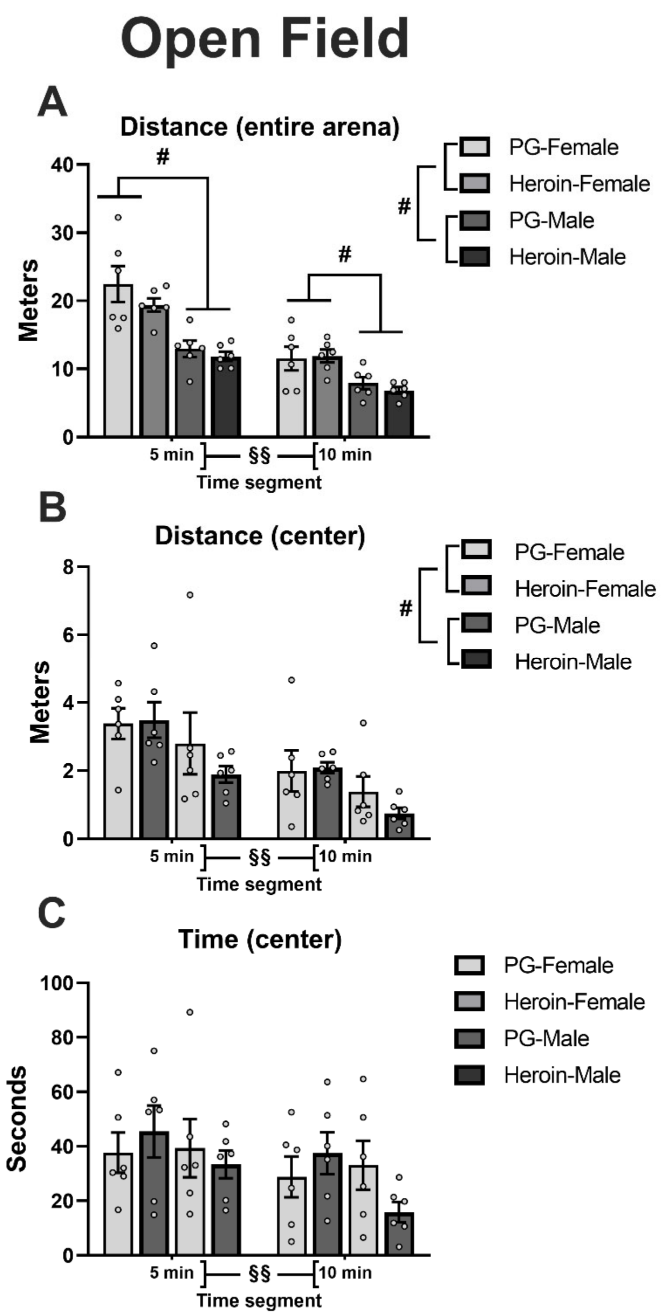


***Fig. S4** Mean (±SEM; N=6 per group) A) total Distance traveled, B) Distance traveled in the Center zone, and C) amount of Time spent in the Center zone in the open field test for female (N = 12) and male (N=12) rats at 15 weeks of age. A significant difference between sexes is depicted with #; a difference between time segments with §§.*

**Open Field**

More distance was traveled by rats in the first five-minute segment compared with the second five-minute segment of the open field test, and female rats traveled significantly more distance compared with male rats (**Fig. S4A)**. The statistical analysis confirmed a significant effect of Time ($F (1, 20) = 299.4$, $p <$ $0.0001$) and of Sex ($F (1, 20) = 24.46$, p

$< 0.0001$) on distance traveled in the entire arena. Additionally, the analysis confirmed significant effects of the interaction of Time with Adolescent Drug treatment ($F (1, 20) = 4.386$, $p < 0.05$) and of Time with Sex ($F (1, 20) = 26.12$, $p <0.0001$). The post-hoc analysis furtherconfirmed that female rats traveled significantly more distance than did male rats in each five-minute bin of the test.  Follow-up analysis of the interaction of Time with Adolescent Drug treatment did not confirm any differences. The analysis of Distance Traveled in the *Center* zone (**Fig. S4B)** confirmed significant effects of Time ($F (1, 20) = 37.88$, $p <0.0001$) and Sex ($F (1, 20) = 5.351$, $p < 0.05$), but no effects of Adolescent Drug. Rats traveled significantly more distance in the Center during the first five minutes of the test than during the last five, and female rats traveled significantly more distance in the Center compared with males. For Time Spent in the Center zone (**Fig. S4C)**, only an effect of Time bin was confirmed ($F (1, 20) = 8.597$, $p < 0.01$).

.
